# Sulci of the canine brain: a review of terminology

**DOI:** 10.1101/374744

**Authors:** Kálmán Czeibert, Patrizia Piotti, Örs Petneházy, Enikő Kubinyi

**Author notes:** Corresponding author: (KC).

## Abstract

Over the last decades there have been several publications of anatomical and neurological textbooks, which include descriptions about the dogs’ brain. However, the terminology used is inconsistent, partly due to individual differences in neocortical gyration and partly due to the common practice of adapting terms from human and murine anatomy. In order to identify such incongruences, in Study 1, we reviewed the existing literature and identified the common terms used as well as any discrepancies between textbooks. Three main forms of inconsistencies were found; a) the use of terms that are not included in the Nomina Anatomica Veterinaria (NAV), b) the inclusion of structures that are listed as not canine-specific, and c) the use of similar names to identify potentially different anatomical structures. To address these issues, in Study 2 we investigated the consistency in appearance of the cerebral sulci, performing a macroscopical examination on 79 canine brains obtained through the Canine Brain and Tissue Bank (CBTB). We then evaluated whether sulci on the frontal regions of brachycephalic breeds differed from those of mesocephalic and dolichocephalic groups, as frontal and olfactory regions are subjected to the most extreme modifications following the shortening of the skull. The statistical analysis showed no difference across the skull length types regarding the occurrence of these sulci, although furrows on the lateral side of the brain proved to be more stable than those on the medial side. In Study 3, we summarized the findings in accordance with the NAV to produce a definitive index of the terms that we recommend be used for each identified sulci. Such an index is beneficial for educational, clinical use, and research (e.g. neuroscience) purposes. The dog is emerging as a pioneering and exceptional model in comparative neuroscience, and therefore the implications for canine neuroscience research should not be underestimated.

## Introduction

Several areas of neuroscience research and clinical activities rely on structural and functional imaging techniques, e.g. magnetic resonance imaging (MRI), computed tomography (CT), positron emission tomography (PET), single photon emission tomography (SPECT), which allow for detailed examination of the central nervous System (CNS). The diagnostic and research interpretation of such techniques requires the knowledge and unambiguous identification of areas and regions; one of the most complex CNS structures being the brain. Brain atlases are therefore fundamental tools for several clinical and research purposes [1–5]. One essential aspect of such research activities is the need to compare, and thus identify, brain structures and areas across different modalities (from macroscopic modalities to histological studies). For this to be done reliably, there is a need for tools such as universal atlases (i.e. an atlas that allows viewing the same structure as depicted through various indirect imaging (CT, MR) or through a more direct approach (surgical intervention and endoscopy)). Consequently, the comparison of brain structures under different imaging modalities requires several atlases, and those currently available in the literature are rarely standardized in their terminology and present a large variability in terminology [6–9]. Thus, the first step to creating a comparative brain atlas for dogs requires the specification of a common descriptive language and clarification of the present terminology in order to make it comparable for every user.

In order to facilitate the establishment of universally accepted anatomical descriptions, the World Association of Veterinary Anatomists (WAVA, http://www.wava-amav.org) regularly provides an internationally recognized list of terms, which describes the macroscopic anatomy of various non-human animals, including dogs. Since 1968, when the first *Nomina Anatomica Veterinaria* (NAV) was published, the International Committee on Veterinary Gross Anatomical Nomenclature (ICVGAN) has regularly updated its recommendation for the use of various anatomical terms. The latest version of NAV available at the time this manuscript was written was the recently revised 6th edition, which was compiled by the ICVGAN, authorized by the General Assembly of the World Association of Veterinary Anatomists, and published by the Editorial Committee [10]. Despite the existence of such guidelines, when describing the morphology of the basic structures of the canine brain (gyri and sulci, i.e. convolutions and involutions visible on the brain surface), there are still some incongruences in the terminology used in veterinary anatomy textbooks (e.g. *Gyrus suprasylvius caudalis* is named *Gyrus ectomarginalis caudalis* [11–14], or *Gyrus ectosagittalis caudalis* [11,14], or *Gyrus suprasylvius posterior* [6,15]). One reason for these incongruences is that in dogs there are no clearly defined borders for individual brain lobes as there are in humans – where, for example, the border between frontal and temporal lobes is defined by a well-recognized anatomical element, the *Sulcus centralis* [16]. Furthermore, the surface morphology of dogs’ brains can vary slightly, even between individuals of the same breed type, due to the different patterns of localization and/or length of their sulci [17,18].

One of the main issues that arises from such nomenclature incongruences is that it makes it difficult to perform comparisons between research findings that use different terms. Furthermore, the assessment of intracranial structures (including anatomical variations and potential abnormalities) requires a common descriptive language between clinicians and researchers. Similar issues have been resolved in the case of another species, the domestic cat, through the production of a tissue probability feline cortical brain map. This map consists of a 3-dimensional cortical template based on cytoarchitectonic and electrophysiological findings, as well as the use of common descriptions regarding surface morphology and terminology [19].

We aimed to review the current situation in the naming of the basic structures of the canine brain, highlighting the discrepancies found in different literature sources, and then, based on a study where we evaluated canine brains, we summarize the sulci that are highly prevalent in dogs. We then present a summarizing table and guideline for use. Such a tool could be beneficial for educational, clinical use, and research (e.g. neuroscience) purposes. The implications for canine neuroscience research should not be underestimated as dogs are considered a good translational model for the study of human neurological and psychiatric diseases [20] and changes in the brain, including for example age related changes [21–23]. The latest human research [24–26] relies heavily on the use of fMRI methods, and in parallel there has also been an increase in the number of non-invasive canine fMRI studies, due to the fact that pet dogs exhibit cooperativeness and trainability, and thus can be trained to lie still and unrestrained in the MRI scan [27–29]. Recent studies have tested the effects of dietary restriction and immunotherapy on aging and aging-associated diseases, and the results highlight the translation value of the dog model for aging and Alzheimer’s disease [30–34]. Different dog brain templates are also developed during the past years to help these fields [35– 37]. These examples show how the dog may become a pioneering and exceptional model in comparative neuroscience.

In order to produce a comprehensive summary of the terms used in canine neuroanatomy, we reviewed the literature searching specifically for canine brain cortical features’ terminology and investigated their consistency in a sample of unprocessed dog brains, in three studies as follows: in Study 1, we aimed to assess the level of consistency among the terms used in the veterinary clinical, educational, and research nomenclature for dog brains. In Study 2, we analysed 79 canine brains looking at the consistency and frequency of occurrence of the sulci that were identified in Study 1. Finally, in Study 3, we summarized a finalised set of terms with color-coded figures congruent with the established official nomenclature, which combines the textbooks reviewed in Study 1, and the findings of the brains analysis of Study 2.

## Materials and methods

### Study 1: Review of the literature

Fourteen veterinary anatomy and neurology textbooks were reviewed, focusing on the terms and definitions used to describe the sulci (grooves or furrows) in the cerebral cortex of the canine brain. For each of these, we described the major analogies and incongruences, and reported and discussed the advantages and limitations that they pose for the field of veterinary medicine and neuroanatomy.

#### Selection and assessment of anatomy textbooks

A literature search was conducted for canine and comparative animal neuroanatomy and neurology textbooks with cortical descriptions, from academic publishers (e.g. Elsevier, Parey, Wiley Blackwell, Springer, and Enke). We included the latest editions available of all the veterinary anatomy books that could be obtained through these sources and were published in the English language, from the year 2000 until the time this study was performed (i.e. between November 2016 and September 2017). For some of the textbooks, the original version (German or French) was used, as no English translation was available. Using this criteria, 14 textbooks were selected [6,8,11–15,38–45]. For each book, the description of the canine telencephalic cortical features was assessed, with a focus on the terms and definitions used to describe sulci. The terms were then compared to those provided by the NAV produced by the World Association of Veterinary Anatomists [14].

#### Furrows

This section includes findings relative to fissures and sulci.

Incongruences emerged in the names used to identify certain structures. Specifically, different names were used to describe the *Fissura pseudosylvia*, i.e. *Fissura sylvii* or *Sulcus pseudosylvius* [44], *Sulcus sylvius* [6,40] and both *Sulcus sylvii* and *Sulcus pseudosylvius* [15]. The *Sulcus corporis callosi* was named *Sulcus callosus* in three textbooks [8,42,43]. Additionally, in some textbooks, the suffix "lateralis" was used instead of "marginalis": this resulted in systematic incongruences, such as the use of *Sulcus ectolateralis* instead of *Sulcus ectomarginalis* [6,15], *Sulcus entolateralis* instead of *Sulcus endomarginalis* [15], and *Sulcus lateralis* instead of *Sulcus marginalis* [6,15]. Furthermore, the *Sulcus rhinalis* was referred to as *Fissura rhinalis* [6,44], and the *Sulcus suprasplenialis* as *Sulcus ectosplenialis* [39]. Finally, some textbooks used the more human-related nomenclature based on anterior-posterior terms in place of rostralis-caudalis [6,15].

We identified terms that were commonly used in the textbooks and were not present in the NAV; these were the *Sulcus ectogenualis* [8,13,15,38,42,44] and the *Sulcus ectosylvius medius* [6,15,38,42]. Furthermore, we observed a category of sulci that seldom occurred in certain textbooks. These are for example the *Sulcus rostralis*, *Sulcus posticus*, *Sulcus occipitalis inferior*, *Sulcus lateromedialis superior, Sulcus lateromedialis inferior* and *Sulcus transsecans* [15]. In one textbook, the rostral part of the *Sulcus splenialis*, just before the *Sulcus cruciatus*, was named *Sulcus cinguli* [38]. A groove rostral to the *Lamina terminalis* was mentioned as *Sulcus parolfactorius* [42], and another between the *Corpus callosum* and the *Sulcus splenialis* was referred to as *Sulcus infrasplenialis* [11]. Finally, a number of structures identified by the NAV as specific to ungulates was used to describe specific structures in the canine brain in some of the textbooks. Examples of this were the *Sulcus calcarinus* [13,44], *Sulcus diagonalis* [15], and the *Sulcus proreus* [12,15,38,42].

A different type of incongruence emerged regarding the identification of certain sulci. We identified inconsistencies in relation to the localization of the *Sulcus ansatus* [46]. The following alternatives were found: some described it as a medial continuation from the *Sulcus coronalis* to the *Sulcus marginalis* [14,15,43], or as a rostro-medial branch derived from the junction of the *Sulcus marginalis* and the *Sulcus coronalis* [6,12,38,42,45]. The rostro-medial branch was described in one book as *Sulcus acominis* [15], which marks all the radial grooves from the major sulci as impressions of vessels [46]. Another textbook [11] showed a transverse groove across the *Gyrus marginalis*, which was then referred to as *Sulcus transsecans* [15], while the rostro-medial branch was entitled as *Sulcus postcruciatus*. However, the *Sulcus postcruciatus* is defined by the NAV as a shallow groove caudal to the *Sulcus cruciatus*, and it should not have any contact with the *Sulcus marginalis* or the *Sulcus coronalis* [14]. One book indicated the *Sulcus postcruciatus* as *Sulcus ansatus* [8], while another referred to it in the text as rostro-medial branch, and in a figure as *Sulcus ansatus*, localized between the *Sulcus coronalis* and *Sulcus marginalis* [44].

#### Short discussion

The review of the literature showed various inconsistencies in terminologies. There were three main forms of inconsistencies; a) the use of terms that were not included in the NAV, b) the inclusion of structures that were listed as not canine-specific in the NAV, and c) the use of similar names to identify potentially different anatomical structures. In order to address these issues, we investigated a large number of dog brains to evaluate the occurrence of the anatomical structures described in the various textbooks, as well as the NAV.

### Study 2: Anatomical assessment

#### Canine Brain and Tissue Bank

Earlier studies determined that the anatomy of the canine brain varies between breed types [47,48], as for example, in brachycephalic dogs, the rostral cranial fossa is shorter than in dolichocephalic dogs and the ethmoidal fossa is less deep, thus the position of the frontal lobe and the olfactory bulbs in brachycephalic breeds may differ from breeds with other skull types [49]. There is also a difference between the frequency of occurrence of the known shape variations among the sulci [50]. Therefore, we decided to perform an anatomical examination of the sulci of a large sample of dog brains, with the aim to assess the consistency in the identification of dog brain sulci. We hypothesized that a) the sulci would be more prevalent on the outer (lateral) surface than on the inner (medial) surface (i.e. sulci located in the medial brain areas would be more difficult to identify (as they are shallower)), and b) the sulci of the frontal lobe of brachycephalic breeds would be less developed compared to other (longer headed) breeds (i.e. the sulci in the frontal lobes of brachycephalic dogs would be more difficult to identify compared to mesocephalic and dolichocephalic dogs).

All the specimens were obtained from the Canine Brain and Tissue Bank (CBTB) established in the Department of Ethology, Institute of Biology, Eötvös Loránd University, Budapest, Hungary. The CBTB is a collection of biomedical material stored under formalin- and/or cryogenic conditions. The aim of the CBTB is to collect tissue samples for research, educational and collaborative purposes. The tissues were obtained from canine bodies, which were donated by dog owners through local veterinary clinics at the time when the animal had to be euthanized for medical reasons.

#### Ethics statement

The brains were obtained from dogs after they had been euthanized for medical reasons (that had no effect on the central nervous system gross morphology), and their bodies donated for research purposes by the owners, in accordance with the Institutional Animal Welfare Committee’s ethical regulations of the University of Veterinary Medicine, Budapest and the Biology Institution of Eötvös Loránd University, Budapest.

#### Subjects

A total of 158 brain hemispheres (N = 158) were obtained from 79 adult dogs (*Canis familiaris*) over one year of age and were examined post-mortem. Only dogs without gross brain deformities were included in the study. The dogs’ bodies had been used for other research purposes prior to the current study; therefore, information about the sex, exact age, and breed of the dogs could not always be obtained at the time of the brain anatomical assessment. However, the dogs’ head morphology group could be extrapolated from craniometrical data. The sample consisted of 19 brachycephalic dogs (Skull index ± SD = 0.68 ± 0.07, min = 0.60, max = 0.88), 47 mesocephalic (Skull index ± SD = 0.55 ± 0.03, min = 0.50, max = 0.61), and 13 dolichocephalic (Skull index ± SD = 0.47 ± 0.02, min = 0.43, max = 0.49).

#### Methods

##### Dissection procedure and skull measurements

Dissection began with the removal of the skin and the temporal muscles in order to identify the main landmarks for the craniometric analysis (shown in Fig 1). The first two landmarks provided the rostro-cranial delimitation of the skull; they were the most rostral point of the incisive bone (prosthion, i.e. the most rostral point of the incisive bone) and the most caudal point of the skull, i.e. where the *Crista sagittalis externa* meets with the *Crista nuchae* (inion / *Protuberantia occipitalis externa*) [42]. After removing the calvaria, the brain was removed from the neurocranium.

**Fig 1.**
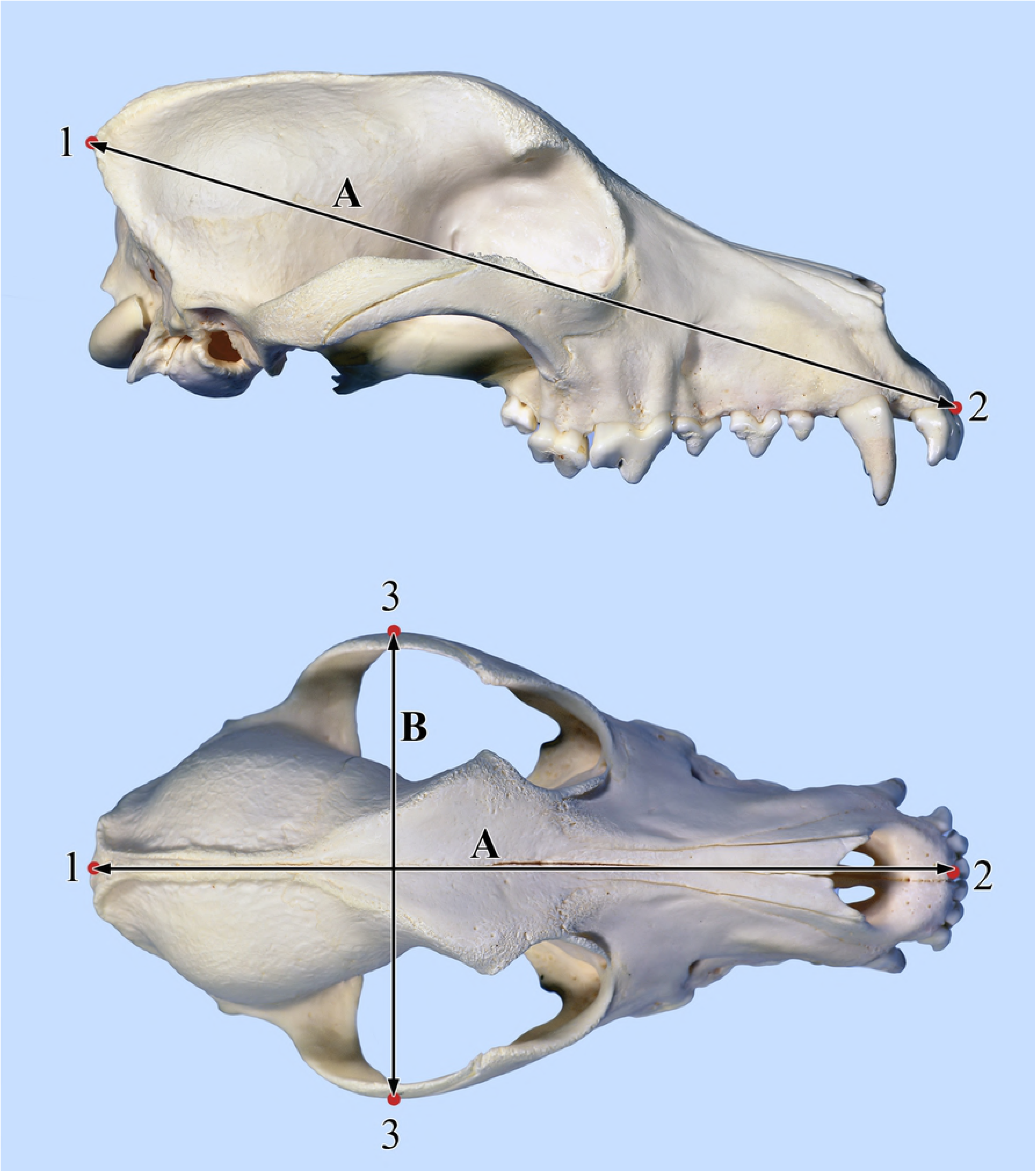
Craniometric indices. 1) inion, 2) prosthion, 3) most lateral point of the zygomatic arch, A) skull length, B) skull width

##### Brain fixation

Brains were initially placed into a rinsing 10% formalin solution for one hour, which ensured the removal of the majority of debris and blood. They were then placed into a 4% formalin solution and conserved for a minimum of two weeks at 4 °C. After this period, the brains were removed from the formalin solution and the two hemispheres were separated with a microtome blade along the midline. The median plane was localized along the *Fissura longitudinalis cerebri*, longitudinally through the *Corpus callosum* and the vermis of the cerebellum, resulting in the separation of the two hemispheres and the two halves of the brainstem.

##### Brain scoring

The sulci were evaluated based on a 4-point scale to assess the presence and ease to recognize each individual sulcus among those identified in Study 1 by one of the authors (KC) who, at the time of assessment, was unaware of the hypothesis of the study. A score of 1 was given if the sulcus was clearly distinguishable (normal), 2 if it was less distinct/shallower than the average or interrupted in multiple sites (less developed), 3 if it was not present (missing), and 4 when it could not be identified (unidentifiable, e.g. due to injury of the affected region).

##### Analysis

Before statistical analysis, a number of indices were calculated to classify dog types. Skull length was defined as the distance between the inion and the prosthion, and the skull width was defined as the distance between the widest points of the zygomatic arches (see Fig 1). The skull index (SI) was then used to identify the head morphology [42] as:

*SI* = *greatest zygomatic width* / *skull length*.

Based on literature recommendations [42], each specimen’s head-type was classified as dolichocephalic (*SI* < 0.5), mesocephalic (0.5 ≤ *SI* < 0.6) or brachycephalic (0.6 ≤ *SI*).

The sulci were grouped in two ways for further analysis. Initially, we identified the sulci located in the medial area of the brain and those located in the lateral area of the brain. Scores of the sulci in the two groups were averaged and compared to assess the consistency in the identification of sulci based on their location. The *sulcus corporis callosi* and the *sulcus hippocampi* were not included in the analysis, because they were not visible without dissecting the brain to seek for hidden deeper structures. We then looked at how certain sulci varied based on the head type. As the rostral part of the brain is likely to vary the most in brachycephalic dogs [49], as compared to dolicho- and mesocephalic dogs, we selected the most rostral sulci for further analysis. The selected sulci were: *Sulcus proreus*, *Sulcus intraproreus*, *Sulcus genualis*, *Sulcus endogenualis*, and *Sulcus rostralis*. Data were analysed using the statistical software IBM SPSS version 22. Kolmogorov-Smirnov tests indicated that some of the variables were not normally distributed, therefore the data were analysed using non-parametric tests. All tests were two-tailed and the level of significance was set at 0.05.

#### Results

The scoring revealed that 31% of the sulci could be identified in more than 90% of the hemispheres and 33% of the sulci were identifiable in less than 10% of the hemispheres (Fig 2). As expected, the *Fissura longitudinalis cerebri* was identified in all hemispheres (as it should separate the two hemispheres, and only in pathological conditions disappears, where the two halves of the brain fuses), also the *Sulcus rhinalis lateralis pars rostralis* could be identified in all dogs, while the *Sulcus rostralis* was missing in the majority of the dogs. In order to avoid pseudoreplication, for each sulcus of each dog, a mean score was calculated for the sulci from the left and right hemispheric scores, which was then used for further analysis.

**Fig 2.**
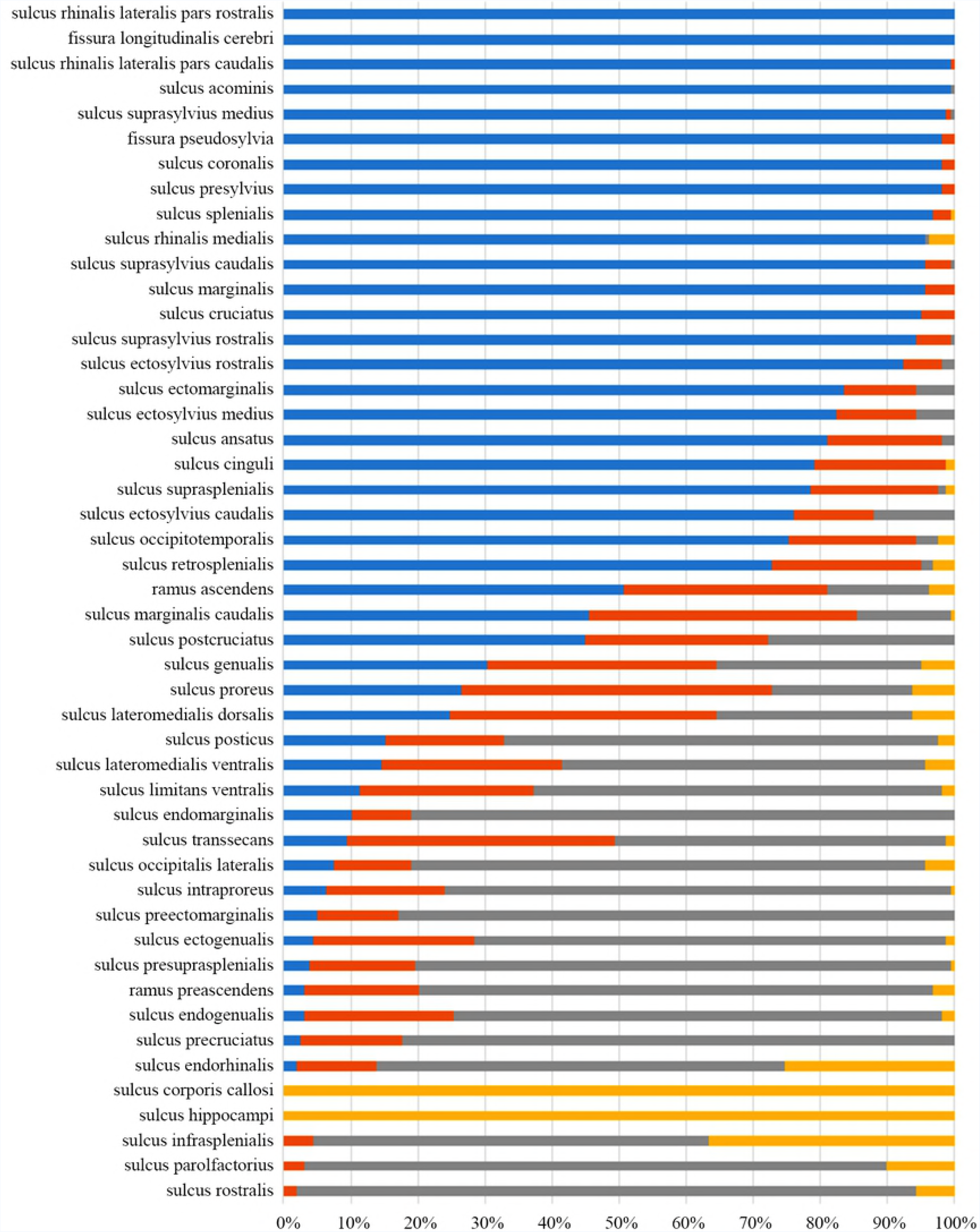
Sulcal frequency of occurrence according to the scoring system. Blue: normal (score 1), red: less developed (score 2), grey: missing (score 3), yellow: unidentifiable (score 4).

The first analysis we performed pertained to the relationship between brain areas and appearance of the sulci. Specifically, we compared the scores of the sulci on the medial part of the brain with those on the lateral part. For each dog, a cumulative score was calculated for the medial area (“medial score” = mean score of all the sulci in the medial area) and for the lateral area (“lateral score” = mean score for all the sulci in the lateral area). A Wilcoxon signed-rank test revealed that the lateral score was significantly lower than the medial score (Mdn_Lat_ = 1.66, Mdn_Med_ = 2.37, z = 7.72, p = 0.001, r = 0.61, Fig 3), thus sulci in the lateral area are more frequently recognizable.

**Fig 3.**
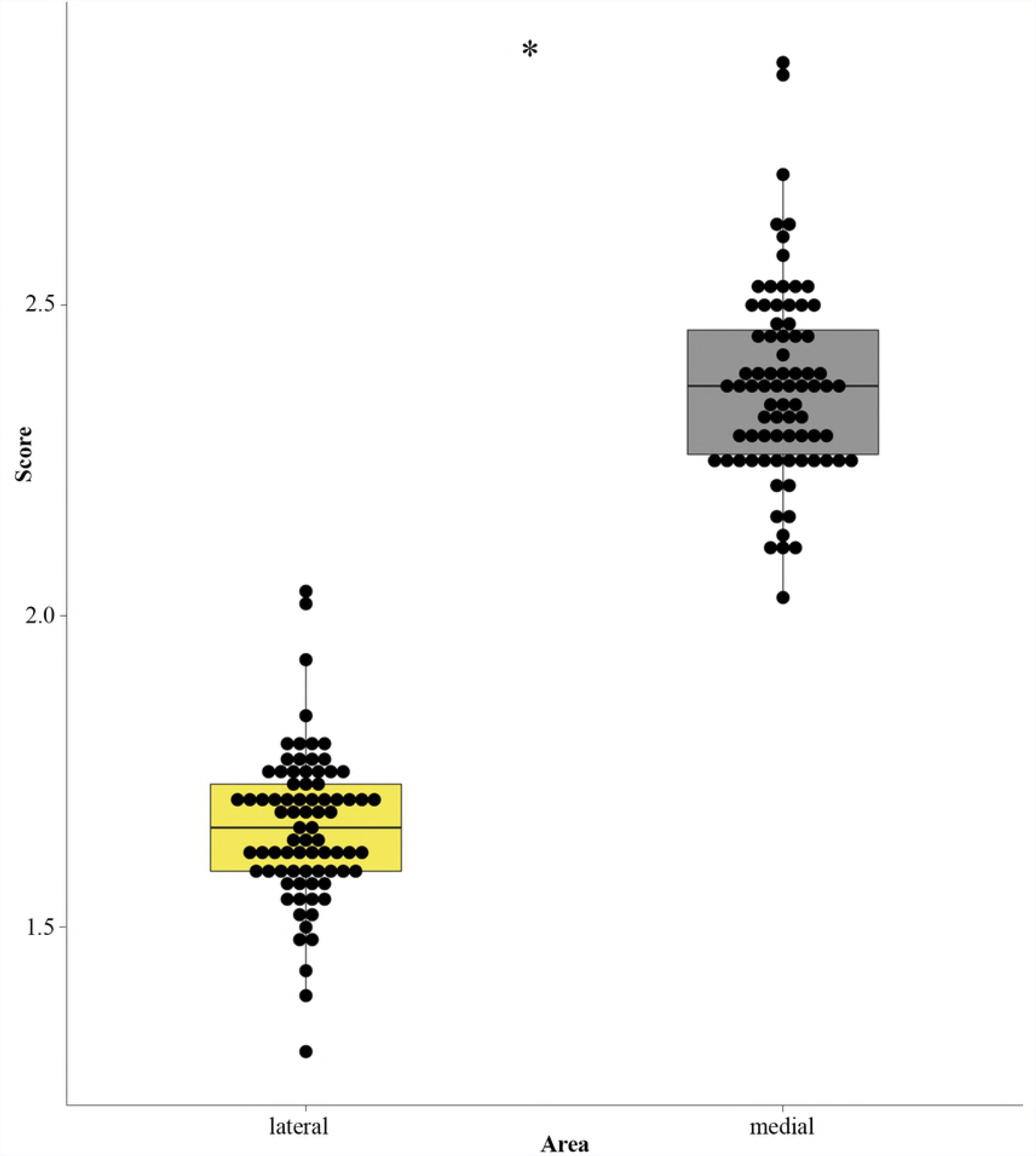
Result of the Wilcoxon signed-rank test. The box-plot represents the cumulative score for each brain area (lateral, in yellow, and medial, in grey). The boxes represent the upper (75% of the scores fall below the upper quartile) and lower (25% of scores fall below the lower quartile) quartiles and the horizontal thick line represents the median. The dots represent the scores of individual dogs (based on the 4-pont scale of assessment). A Wilcoxon signed-rank test indicated a significant difference between the scores of each lateral and medial brain area (* = p < 0.05).

We then investigated the differences in the selected sulci, based on head type. Due to shift in the olfactory bulb and the frontal lobe in brachycephalic breeds compared to other (meso- and dolichocephalic) head types [49], we expected that it could have an effect on the appearance of the sulci at this region. Therefore, we compared the score of the most rostral sulci (both individually and as a group) between brachycephalic dogs and other head types. Mann-Whitney U tests revealed that there were no significant differences between brachycephalic dogs and other head types in any of the individual sulci or the cumulative score (Table 1).

**Table 1.**
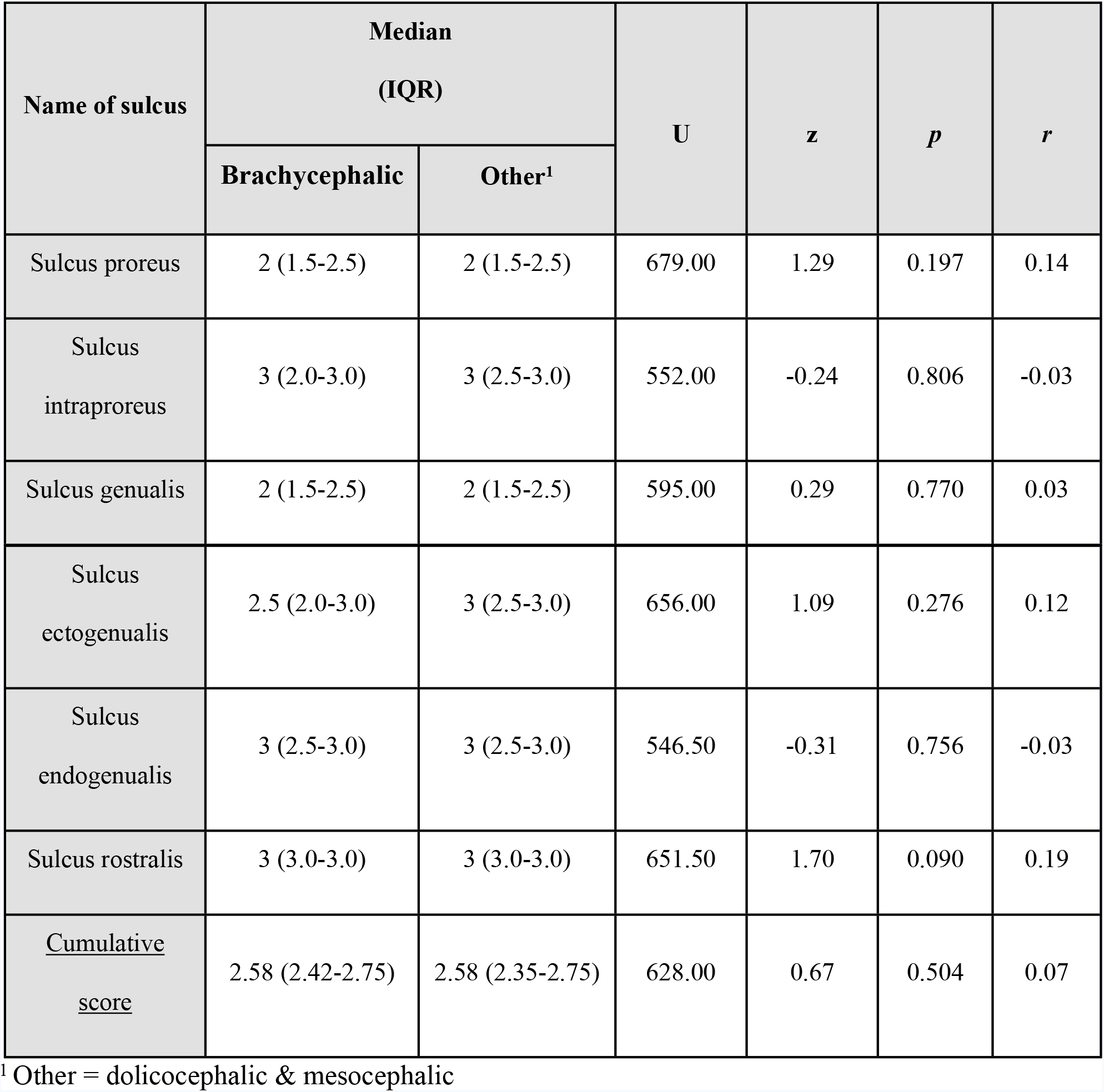
Results of the Mann-Whitney U tests comparing brachycephalic dogs and other head types in the individual sulci and cumulative score. Test statistics are indicated as “U”, standardized test statistics are indicated as “z” and effect size as “*r*”; statistical significance was set at *p* = 0.05.

### Study 3: Terminology summary

Following the results of Study 1 and Study 2, we summarized the index of the canine brain’s main sulci for further discussion and future terminology guidelines. We based the index on the terms that showed the highest consistency across the two modalities of analysis. Thus in Table 2 we provide the terms that primarily according to the NAV’s guidelines but also broadening its spectrum we recommend be used for each identified sulci and compared them to those that have been used in the past, according to the literature reviewed, so that it is possible to identify the term of reference starting from any term used in the literature considered in this work.

**Fig 4.**
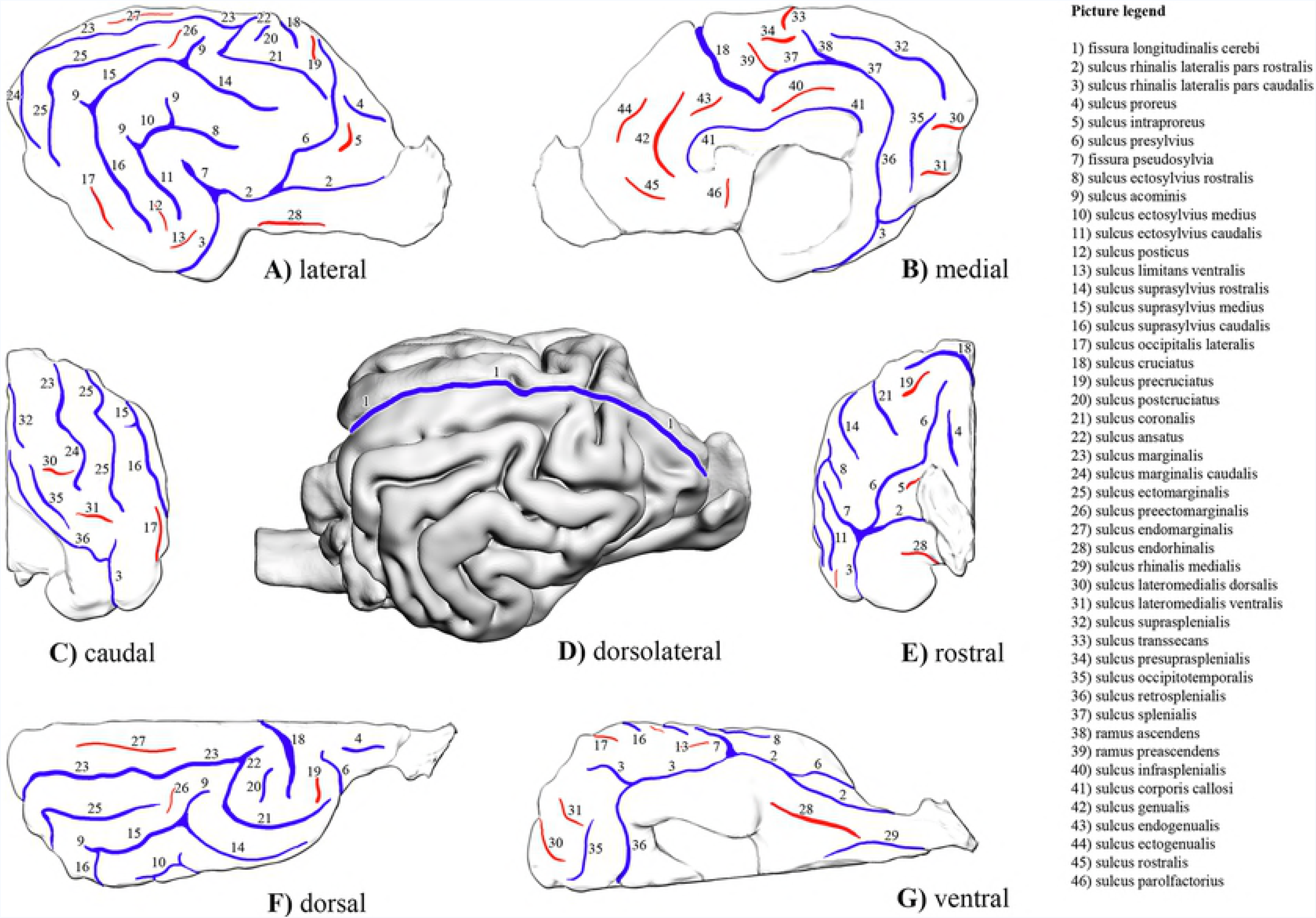
Cortical map of the main sulci of the canine brain. A) Lateral view of the right hemisphere, B) medial view of the right hemisphere, C) caudal view of the right hemisphere, D) dorsolateral view of the entire brain (3-dimensional model), E) rostral view of the right hemisphere, F) dorsal view of the right hemisphere, and G) ventral view of the right hemisphere. Sulci that were recorded as missing or indistinct in less than 33% of the cases in Study 2 are marked with red, and the more prevalent ones (sulci that were distinguishable the most) are marked with blue.

Terms that were consistently used included across textbooks *Fissura longitudinalis cerebri* (Fig4/1), *Sulcus ansatus* (Fig4/22), *Sulcus coronalis* (Fig4/21), *Sulcus endorhinalis* (Fig4/28), *Sulcus genualis* (Fig4/42), *Sulcus hippocampi, Sulcus presylvius* (Fig4/6), *Sulcus rhinalis medialis* (Fig4/29), *Sulcus splenialis* (Fig4/37), *Sulcus ectogenualis* (Fig4/44) and *Sulcus precruciatus* (Fig4/19). For the terms that had more than one alternative name based on the literature, we recommend the use of the following terms (see also Table 2): *Fissura pseudosylvia* (Fig4/7), *Sulcus corporis callosi* (Fig4/41), *Sulcus cruciatus* (Fig4/18), *Sulcus ectomarginalis* (Fig4/25), *Sulcus ectosylvius caudalis* (Fig4/11), *Sulcus ectosylvius rostralis* (Fig4/8), *Sulcus endomarginalis* (Fig4/27), *Sulcus marginalis* (Fig4/23), *Sulcus postcruciatus* (Fig4/20), *Sulcus proreus* (Fig4/4), *Sulcus rhinalis lateralis pars rostralis* (Fig4/2), *Sulcus rhinalis lateralis pars caudalis* (Fig4/3), *Sulcus suprasplenialis* (Fig4/32), *Sulcus suprasylvius caudalis* (Fig4/16), *Sulcus suprasylvius medius* (Fig4/15), *Sulcus suprasylvius rostralis* (Fig4/14), *Sulcus ectosylvius medius* (Fig4/10), *Sulcus endogenualis* (Fig4/43), *Sulcus intraproreus* (Fig4/5), *Sulcus retrosplenialis* (Fig4/36),and *Sulcus marginalis caudalis* (Fig4/24).

For those terms that were mentioned in only one source, we recommend: *Sulcus lateromedialis dorsalis* (Fig4/30), *Sulcus lateromedialis ventralis* (Fig4/31), *Sulcus limitans ventralis* (Fig4/13), *Sulcus occipitalis lateralis* (Fig4/17), *Sulcus occipitotemporalis* (Fig4/35), *Sulcus posticus* (Fig4/12), *Ramus ascendens sulci splenialis* (Fig4/38), *Sulcus preectomarginalis* (Fig4/26), *Sulcus presuprasplenialis* (Fig4/34), *Sulcus rostralis* (Fig4/45), *Sulcus transsecans* (Fig4/33), *processus acominis* (Fig4/9), *Sulcus infrasplenialis* (Fig4/40), *Sulcus parolfactorius* (Fig4/46). As the *Ramus ascendens sulci splenialis* (38) and the *Sulcus occipitotemporalis* (Fig4/35) were found in more than 33% of the dogs in Study 2, and their presence can serve as a good landmark for MRI analysis or surgical interventions, we recommend their use even though at the moment they are mentioned only in one source [15]. Furthermore, radially directed grooves, which extend from the ectosylvian and suprasylvian grooves into the dorsal direction and have regularly blood vessels crossing on them, should be called *Sulci acominis* based on the recommendation of some authors [15,46].

**Table 2.**
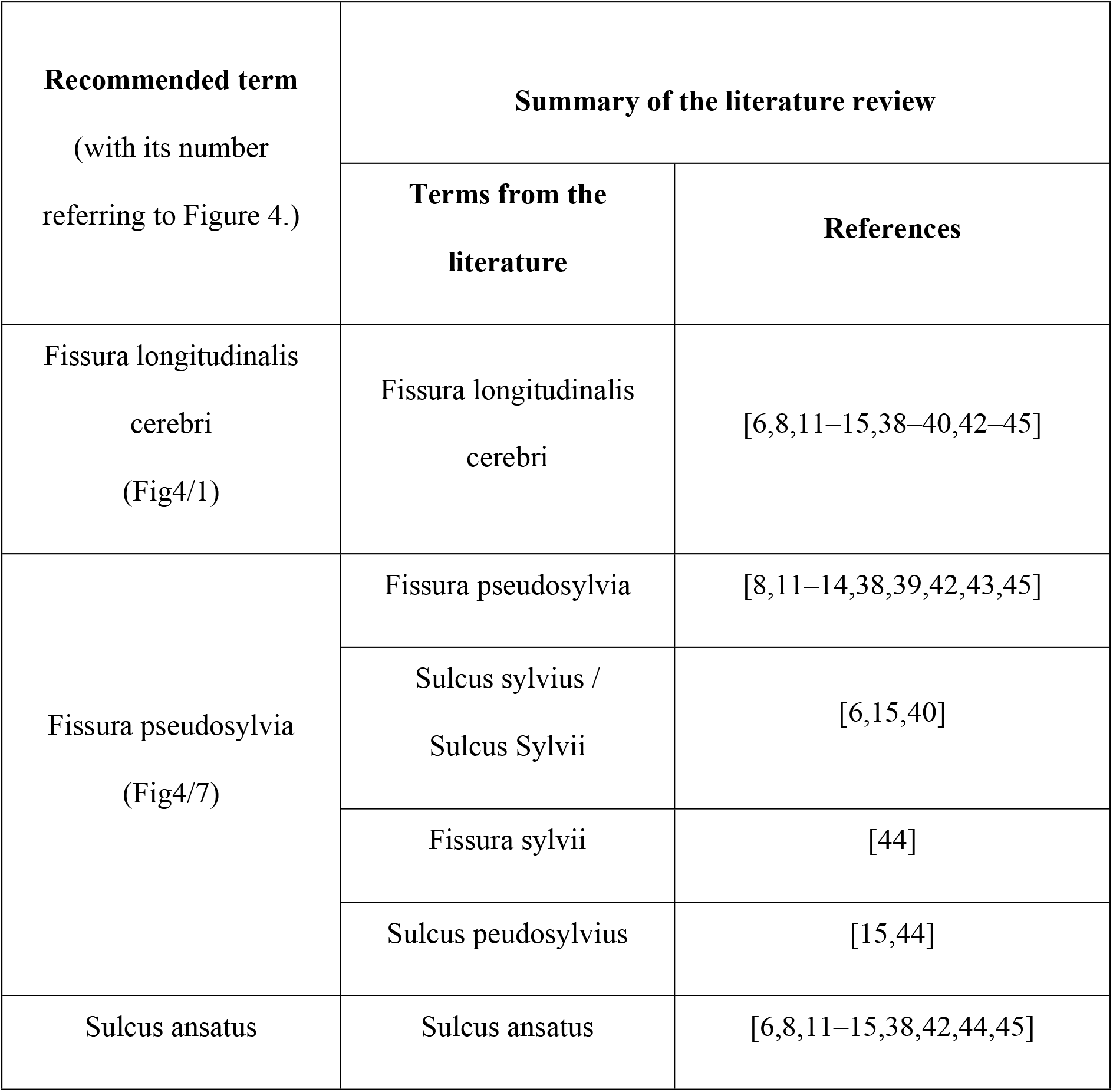
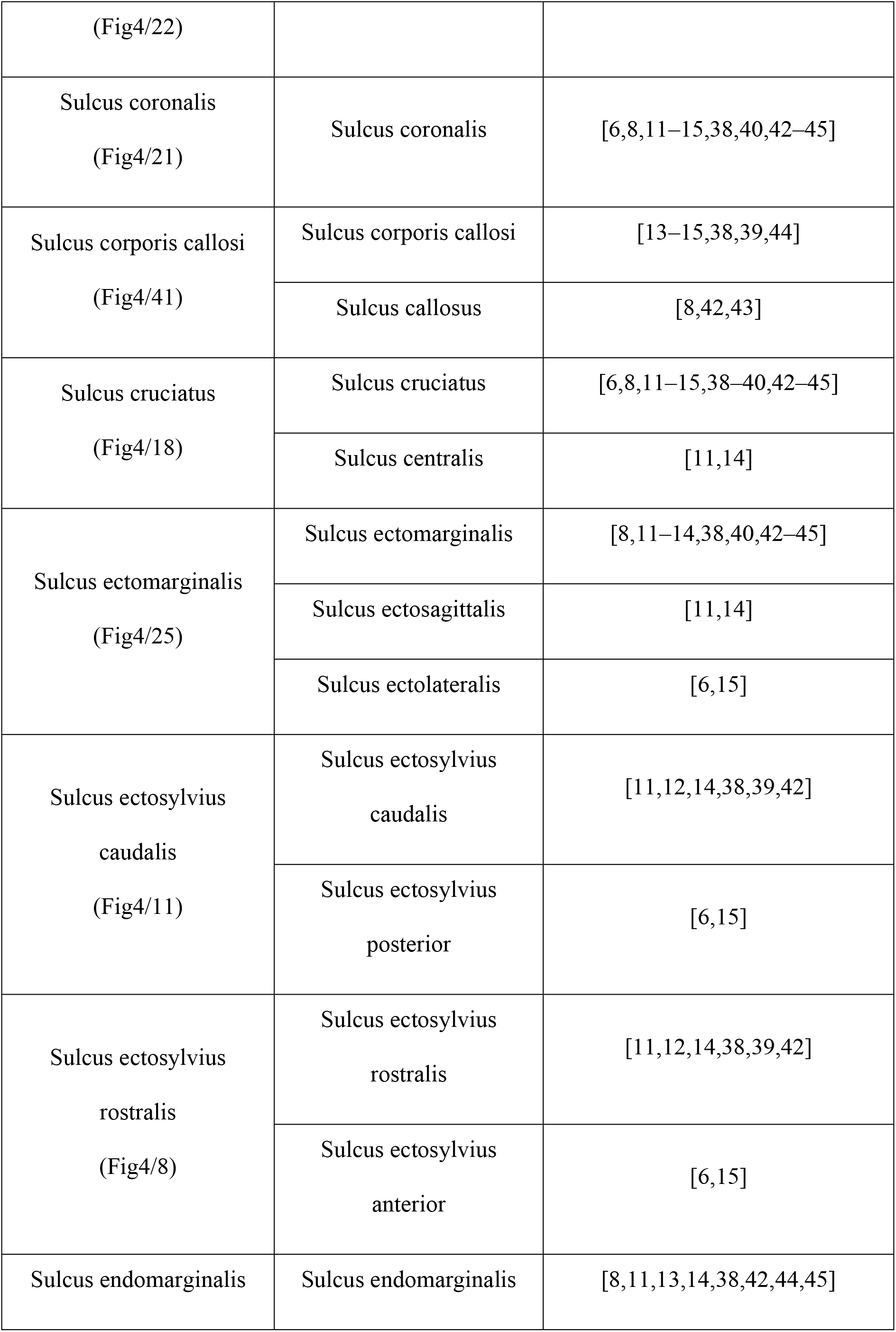
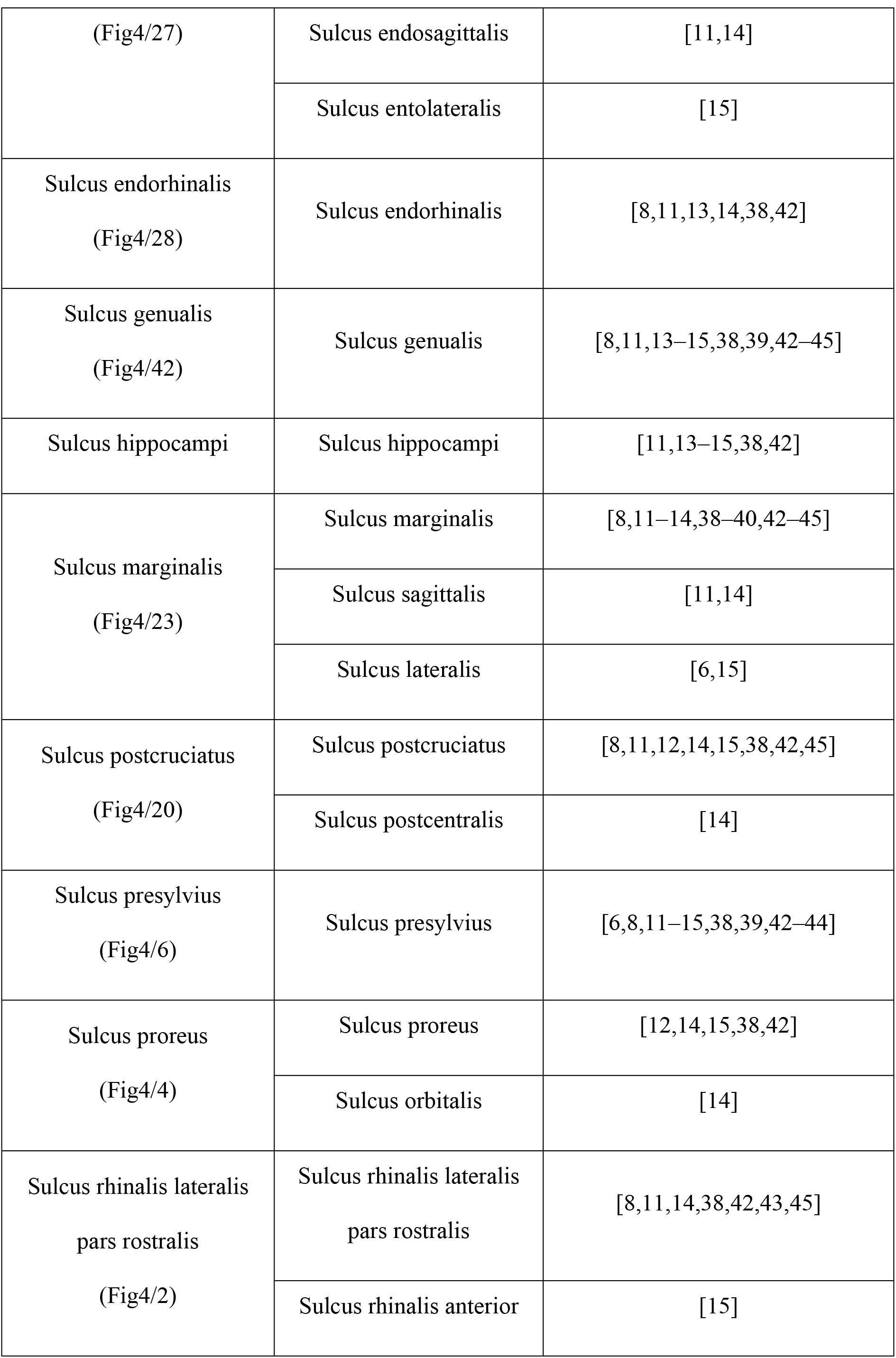
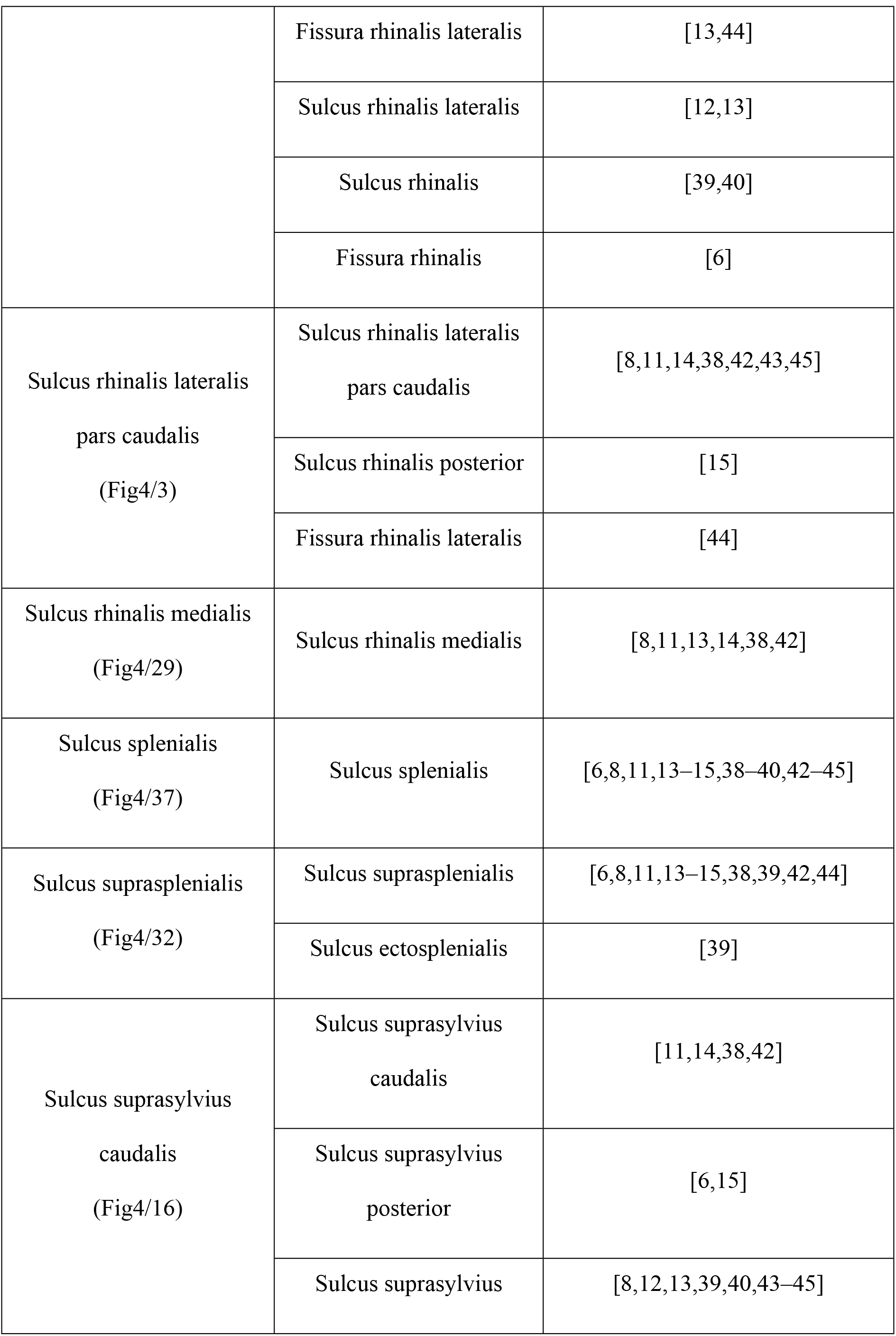
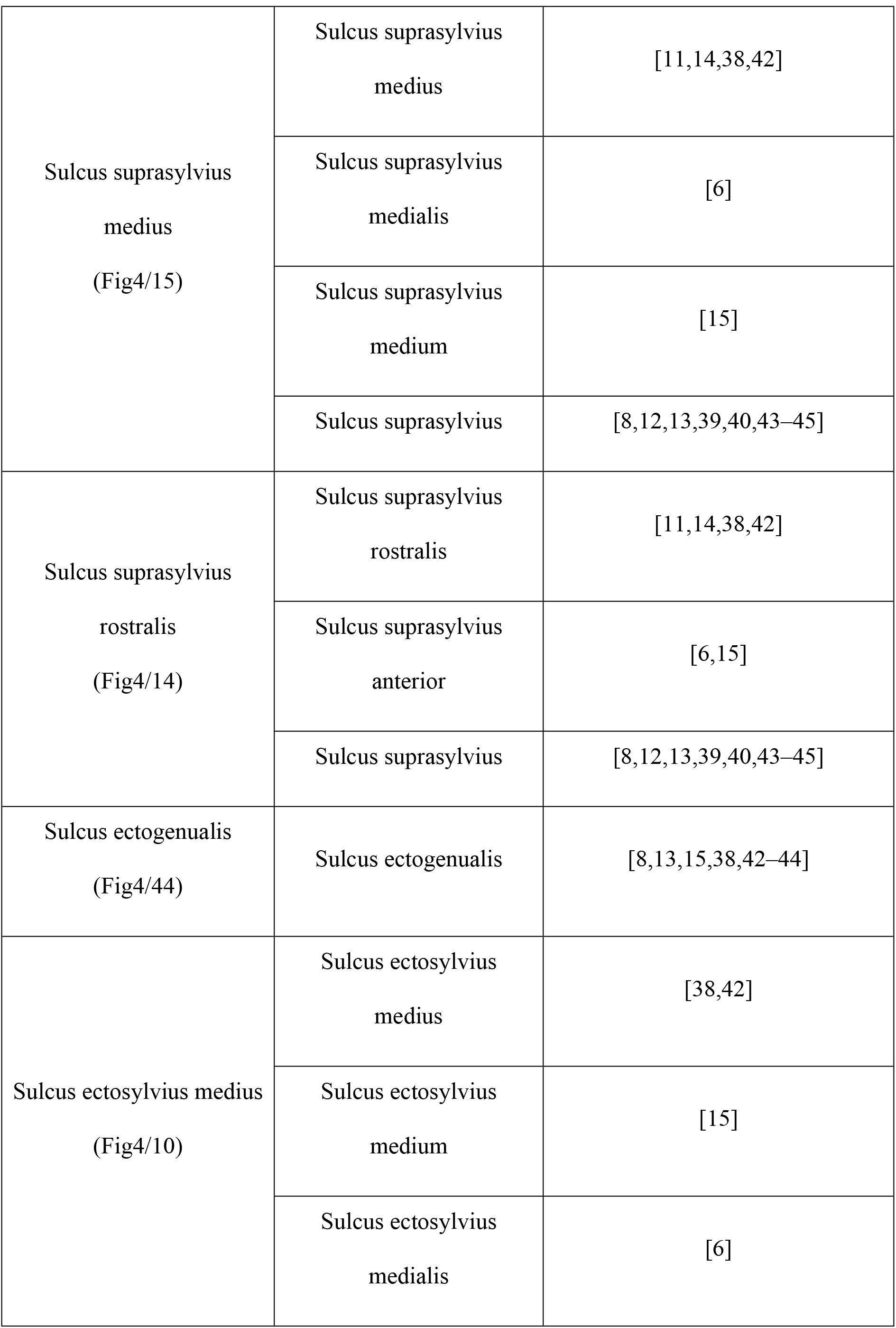
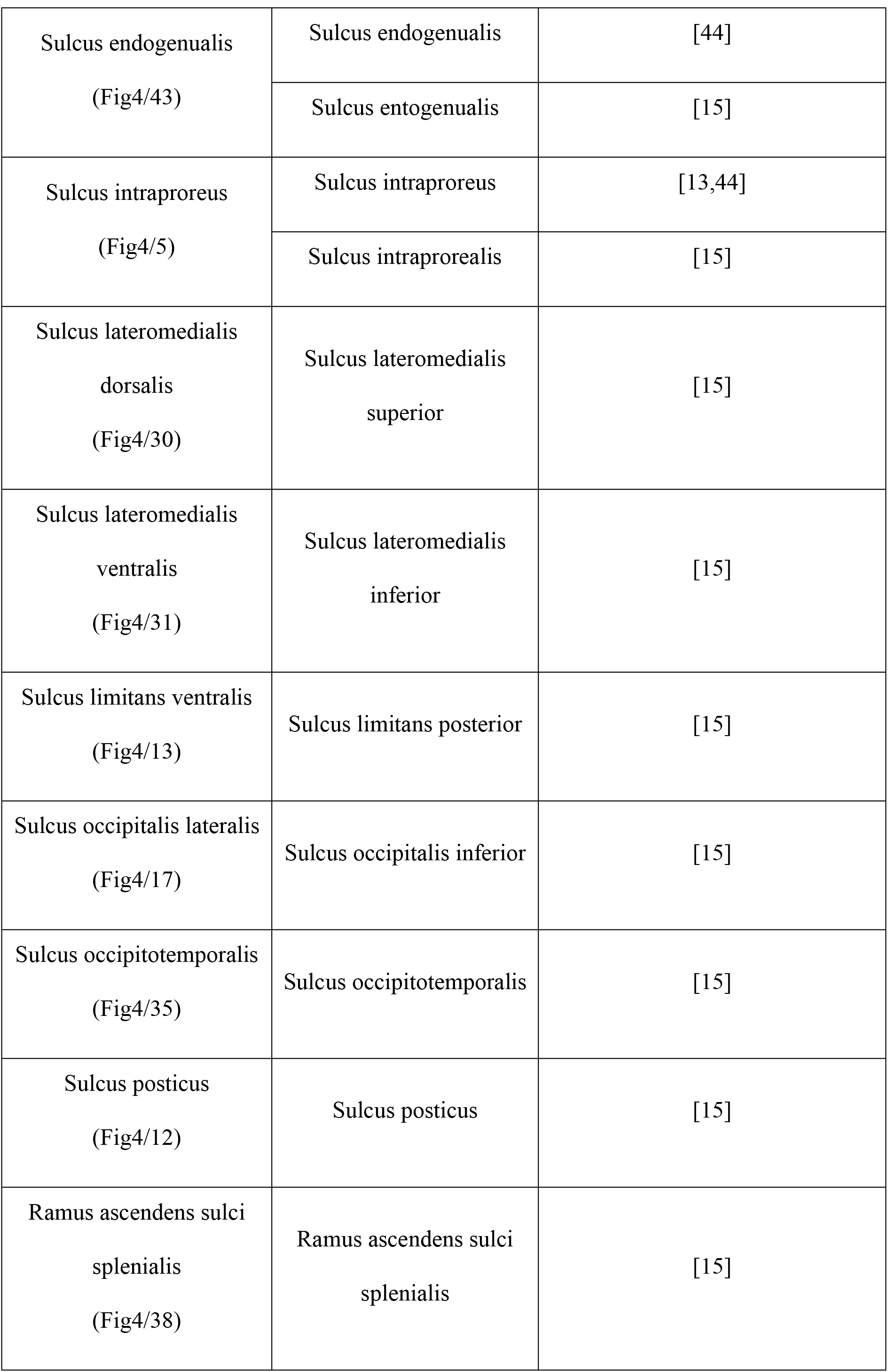
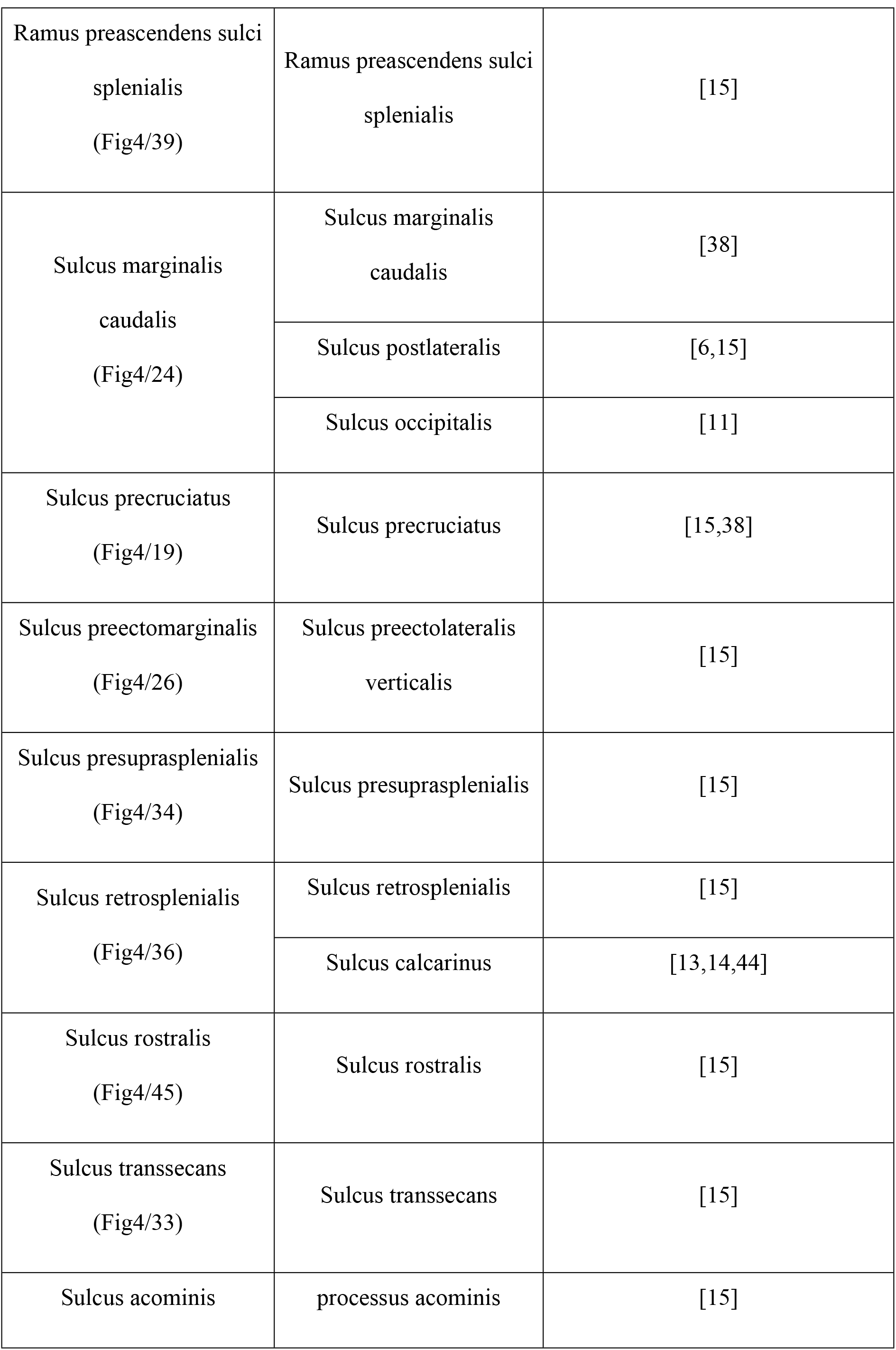
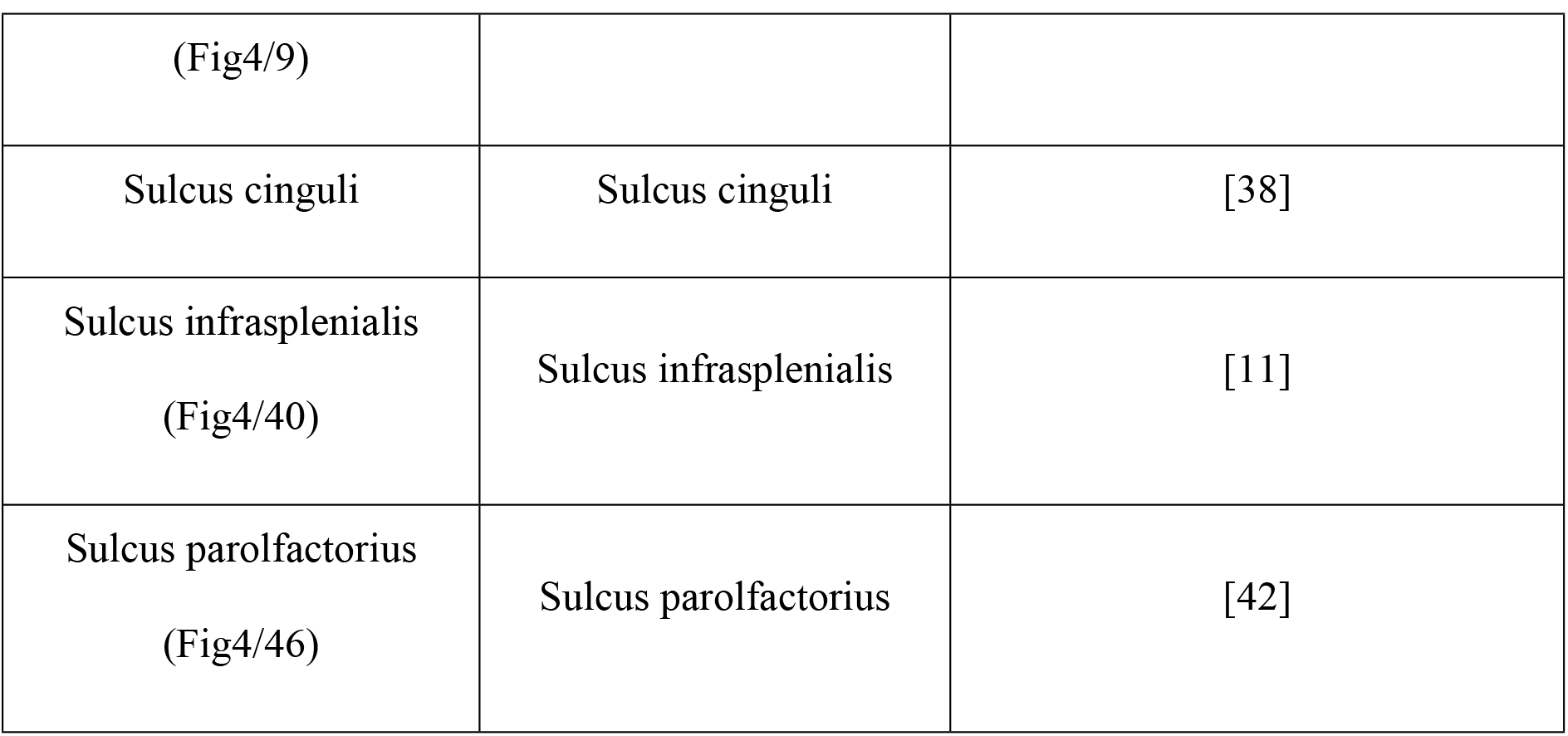
Index of sulci. The index shows the terms used in the literature to describe the sulci of the canine brain and the proposed terms.

## General discussion

We reviewed the commonly used terminology regarding the sulci of the canine brain in order to assess the occurrence of possible inconsistencies. We often found several different anatomical terms that refer to the same cortical structures, which can cause difficulties in interpreting comparative studies.

To our knowledge, this is the first publication to compare the anatomical description of canine sulci used in the literature. Although the Illustrated Veterinary Anatomical Nomenclature, based on the Nomina Anatomica Veterinaria (NAV) [14], provides a reliable official collection of recommended terms for the description of veterinary anatomy with figures, the use of such terms, as found in the current study, is not always widespread across brain anatomy textbooks. A review of academic textbooks was performed in Study 1, which compared fourteen major books in the field of comparative anatomy, neuroanatomy and neurology, to reveal differences among the sources in the terminology used to label the different cortical structures. We showed that several terms have been derived from human terminology, especially the directions (anterior/posterior, superior/inferior), and are used in the veterinary literature. We recommend the use of the veterinary-related terms because most of the intracranial structures (vessels, nerves, and pathways) are named according to the NAV, using dorsal/ventral instead of superior/inferior and rostral/caudal instead of anterior/posterior (e.g. *Sinus sagittalis dorsalis, Nuclei rostrales thalami, Nucleus cochlearis dorsalis et ventralis, Tractus spinocerebellaris dorsalis et ventralis* etc.) [14].

In Study 2, we assessed the presence of the main sulci in a large sample of dogs and we evaluated whether an individual structure was present, underdeveloped, missing or unidentifiable. Using craniometric indexes, we compared whether sulci of brachycephalic breeds differed from those of mesocephalic and dolichocephalic groups, based on the presence of certain sulci on the frontal regions, as this part of the brain is supposed to be subjected to the most extreme modifications following the shortening of the skull. The analysis showed no difference across the skull length types regarding the occurrence of these sulci, although furrows on the lateral side of the brain proved to be more stable than those on the medial side.

One limitation of the evaluation performed in Study 2, is that fixation in formalin *ex situ* can cause flattening on the surface if the brain is not placed properly in the container, and surface blood vessels can make impression on the cortex that appear similar to the original grooves. While knowledge of basic topography may aid in differentiating in these cases, it was not possible to exclude the chance that these types of events occurred. In future studies, the true depth of the sulci would be assessed through T2-weighted MR imaging, where the *Liquor cerebrospinalis* could clearly highlight the grooves. By making 3-dimensional models from the MR images one could also interactively visualise not only the grooves but the gyri as well.

We have provided a thorough review of the current state of the terminology used to describe the canine brain. Our summary and recommendations for appropriate terminology can serve as a base for future canine brain atlases and medical terminology.

## Acknowledgements

We are grateful for Dr. Judit Benczik for commenting on an earlier version of this manuscript. We also would like to thank Professor Dr. Hagen Gasse for his helpful comments and suggestions and to Dr. Lisa Wallis for the language proofing.

